# High interaction valency ensures cohesion and persistence of a microtubule +TIP body at the plus-end of a single specialized microtubule in yeast

**DOI:** 10.1101/2021.09.13.460064

**Authors:** Sandro M. Meier, Ana-Maria Farcas, Anil Kumar, Mahdiye Ijavi, Robert T. Bill, Jörg Stelling, Eric Dufresne, Michel O. Steinmetz, Yves Barral

**Affiliations:** Institute of Biochemistry, Department of Biology, ETH Zürich, Zürich, Switzerland; Paul Scherrer Institute, Villigen, Switzerland; Department of Materials, ETH Zürich, Switzerland; Department of Molecular Life Sciences, UZH, Switzerland; Department of Biosystems Science and Engineering, ETH Zürich, Switzerland; University of Basel, Biozentrum, CH-4056 Basel, Switzerland

## Abstract

Microtubule plus-end tracking proteins (+TIPs) control microtubule specialization and are as such essential notably during eukaryotic cell division. Here, we investigated interactions and functions of the budding yeast Kar9 network consisting of the core +TIPs components Kar9 (functional homologue of APC, MACF, and SLAIN), Bim1 (orthologue of EB1), and Bik1 (orthologue of CLIP-170). Our data indicate that a redundant, multivalent web of interactions links the three +TIPs together to form a “Kar9 body” at the tip of a single cytoplasmic microtubule. They further suggest that this body is a liquid-like condensate, allowing it to persist on both growing and shrinking microtubule tips, and functions as a mechanical coupling device between microtubules and actin cables during mitosis. Our study underlines the power of dissecting the web of low-affinity interactions driving liquid-liquid phase separation of proteins in order to demonstrate the importance and establish the functional roles of condensation processes in vivo.

## Introduction

Microtubules play a central role in moving and positioning diverse cargos within eukaryotic cells and thus in controlling cell physiology. To fulfil these functions, individual microtubules acquire specialized behaviors to carry out dedicated roles independently of each other. For example, kinetochore, interpolar, and astral microtubules of the mitotic spindle show distinct dynamics and functions throughout mitosis (reviewed in (Prosser and Pelletier, 2017)). While in anaphase, kinetochore microtubules shrink to pull sister chromatids apart, interpolar microtubules grow and elongate the spindle. Concomitantly, aster microtubules interact dynamically with the cell cortex to position the spindle and the cleavage apparatus properly relative to each other. To achieve their individual functions, each of these microtubules must have the ability to attach to selected targets, transport specific cargos, and adopt specific dynamics. How microtubules achieve such specialization despite being all composed of the same building blocks is a fundamental open question.

One large class of regulators that contribute to microtubule specialization are the microtubule plus-end tracking proteins (+TIPs). They can be broadly sorted into two groups. The first group comprise ubiquitously expressed and evolutionary conserved proteins that bind the plus-ends of most if not all microtubules (reviewed in (Akhmanova and Steinmetz, 2008, 2015)). These include the end binding proteins (EBs, e.g., EB1, EB2 and EB3 in mammals), which directly bind microtubule tips, and the cytoplasmic linker proteins (CLIPs, e.g., CLIP-170 in mammals), which bind microtubules through EBs. These +TIPs typically track growing but not shrinking microtubule tips. The second group of +TIPs, which we called “patterning +TIPs” (Kumar et al., 2021), comprises protein families of diverse phylogenetic origins that typically associate with specific microtubules. Representatives include the microtubule-actin crosslinking factor (MACF/ACF7/Shot, involved in axon guidance (Sonnenberg and Liem, 2007)), SLAIN motif-containing proteins 1 and 2 (implicated in axon growth (van der Vaart et al., 2012)), and the adenomatous polyposis coli (APC, a tumor suppressor acting in cell division (reviewed in (Nelson and Näthke, 2013)). These +TIPs bring together the plus-ends of specialized microtubules with target structures such as the actin cytoskeleton, anchorage sites at the plasma membrane, or chromosomes (reviewed in (Akhmanova and Steinmetz, 2008, 2015)). How patterning +TIPs, which typically interact with microtubules in an EB-dependent manner, are targeted to and remain associated with dedicated microtubule ends is poorly understood.

The budding yeast *Saccharomyces cerevisiae* offers a powerful model for studying how microtubules become specialized. During mitosis, every cell comprises two spindle pole bodies (SPBs, equivalent of the metazoan centrosomes): an old SPB, inherited from the previous mitosis, and a newly synthesized one (reviewed in (Lengefeld, 2018 #61)). Each SPB nucleates a small aster composed of one to three cytoplasmic microtubules (Gupta et al., 2002; Saunders et al., 1997). Interestingly, these cytoplasmic microtubules are specialized in an SPB-dependent manner. The old SPB forms long microtubules that interact with the cortex of the bud, the future daughter cell. In contrast, the microtubules generated from the new SPB remain highly dynamic and short (Chen et al., 2019; Lengefeld et al., 2018; Segal and Bloom, 2001; Shaw et al., 1997). The interaction of microtubules emanating from the old SPB with the bud cortex ensures that only this single SPB orients towards the bud (Hotz et al., 2012; Pereira et al., 2001). This specialization ensures that the intranuclear spindle aligns properly with the mother-bud axis (the future axis of cell division) before anaphase onset, allowing the elongating spindle to equally distribute the genetic material between the mother and its bud during anaphase (Kusch et al., 2003; Kusch et al., 2002; Liakopoulos et al., 2003).

Specialization of microtubules emanating from the old SPB relies in particular on their exclusive decoration with the protein Kar9, a patterning +TIP that is functionally related to MACF/ACF7/Shot, SLAIN, and APC (Kumar et al., 2021). At the plus-end of one microtubule, which Kar9 binds through the EB-family member Bim1, Kar9 interacts with the CLIP-family member Bik1 and the myosin V actin-directed motor Myo2 (Korinek et al., 2000; Kumar et al., 2021; Manatschal et al., 2016; Miller et al., 2000; Moore et al., 2006). Myo2 then pulls the microtubule plus-end along actin cables towards the bud cortex to drive the specific interaction of the selected microtubule with the bud cortex (Beach et al., 2000; Yin et al., 2000). A unique and important feature allowing Kar9 to remain associated with its target microtubule throughout mitosis is its ability to track its plus-end throughout phases of both microtubule growth and shrinkage (Kusch et al., 2002; Liakopoulos et al., 2003).

The molecular mechanisms underlying Kar9 recruitment to and tracking of the growing and shrinking plus-end of a single, dedicated microtubule in vivo have remained very puzzling. In order to address the fundamental issue of microtubule specialization, we investigated here the mechanisms of Kar9 cohesion and persistence to essentially one microtubule tip in vivo through dissecting the network of interactions that Kar9 establishes with its key partners Bim1 and Bik1.

## Results

### The core components of the Kar9-network undergo liquid-liquid phase separation in vitro

To characterize the interactions of the core components of the Kar9 network with each other, we recombinantly produced Kar9, Bim1, and Bik1 (Figure 1A), and monitored their interactions in vitro using a pelleting assay. Mixed together, the three proteins were soluble in a standard, crowding-agent free, and pH neutral buffer containing 500 mM sodium chloride, but pelleted readily upon ultracentrifugation when the salt concentration was reduced to 200 mM (Figure 1B). Microscopy imaging indicated that under this condition the proteins phase separated into liquid-like droplets (Figure 1C). Removal of Bim1 or Bik1 from the mixture changed the appearance of the droplets, but did not affect coalescence. Remarkably, removing Kar9 fully solubilized the Bim1+Bik1 mixture even at low salt. This was because although Bik1 condensates on its own at low salt, adding Bim1 resolubilized the protein. When alone, Kar9 was soluble only at high salt. However, instead of condensing into liquid-like droplets it collapsed into amorphous aggregates at low salt. Together, these data indicate that while insoluble on its own at low salt, Kar9 promotes the liquid-liquid phase separation of Bim1 and Bik1 in vitro and is itself solubilized in these droplets.

**Figure 1:**
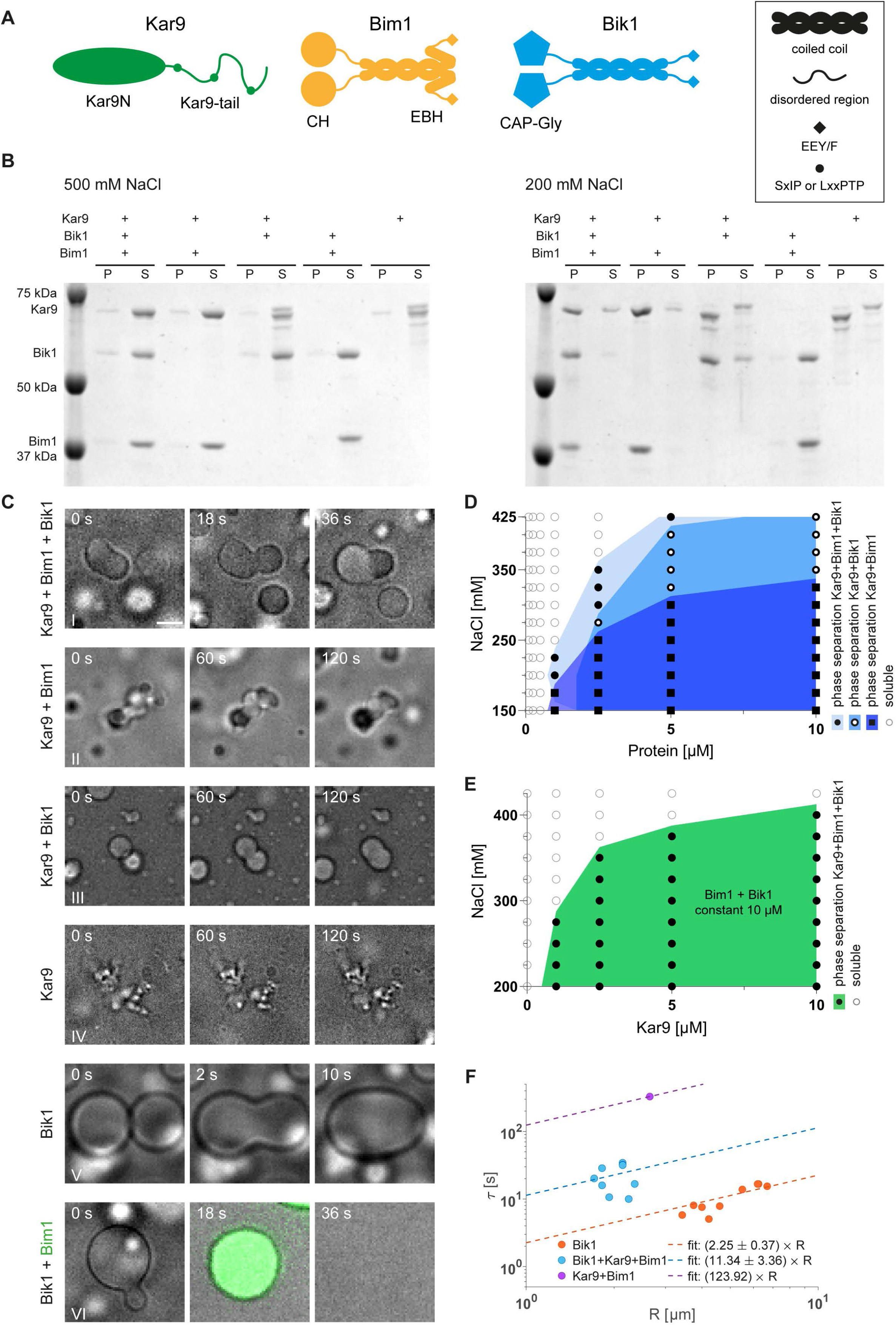
Characterization of Kar9, Bim1, and Bik1 phase separation in vitro. **A**, Schematic representations of Kar9, Bim1, and Bik1. **B**, Pelleting assays of Kar9 and mixtures with Bim1 and Bik1 (10 μM each) analyzed by Coomassie stained SDS-PAGE at 500 (left panel) or 200 mM (right panel) NaCl. P, pellet; S, supernatant. **C**, Time-lapse phase contrast microscopy images of mixtures of Kar9 with Bim1 and/or Bik1 (I-III), Kar9 alone (IV), and Bik1 alone (V) at 10 μM protein and at 220 mM NaCl concentration. For the mixture Bik1+Bim1 (VI), Bik1 droplets were preformed and then Atto-488-NTA-labelled Bim1 was added. Scale bar, 3 μm. **D**, Superimposed phase diagrams of equimolar mixtures of Kar9+Bim1+Bik1 (light blue), Kar9+Bik1 (intermediate blue) and Kar9+Bim1 (dark blue) obtained at different protein and NaCl concentrations. Symbols indicate the presence of droplets; colored areas are the inferred regions where phase separation occurs. **E**, Phase diagram of Kar9 that was titrated into a constant equimolar mixture of Bim1 and Bik1 (10 μM each). **F**, Quantification of the droplet fusion time, τ, as a function of droplet radius, R, based on phasecontrast microscopy movies as shown in (C), presented on a log-log plot. Dots indicate measurements; lines indicate linear fits to the data with reported slopes of τ as a function of R.

To better characterize the phase separation behavior of the Kar9+Bim1+Bik1 mixture, we analyzed its behavior with varying salt and protein concentrations (from 150 to 425 mM NaCl and 0.1 to 10 μM for each protein). Droplet formation was screened by microscopy (Figure 1D). When mixed in equimolar amounts, the Kar9+Bim1+Bik1 droplets remain resistant even towards high salt concentrations. Removing Bik1 caused the Kar9+Bim1 droplets to dissolve above 300 mM salt, and removing Bim1 increased the Kar9+Bik1 mixture’s critical concentration for droplet formation at most salt concentrations tested. At 200 mM salt, the critical protein concentration was 0.5-1.0 μM for Kar9+Bim1+Bik1 and 1.0-2.5 μM for Kar9+Bim1 or Kar9+Bik1. Varying the concentration of solely Kar9 indicated that 1.0 μM of Kar9 was sufficient to induce droplet formation in an equimolar mixture of 10 μM each of Bim1 and Bik1 (Figure 1E). Thus, all three components and particularly Kar9 promoted phase separation of the core Kar9-network components over a broad range of conditions.

Analysis of the size, morphology, and dynamics of the droplets within minutes of dropping the salt concentration indicated that those formed of Bik1 alone were the most fluid, fusing and relaxing within seconds, followed by Kar9+Bim1+Bik1 (tens of seconds), and Kar9+Bik1 and Kar9+Bim1 droplets (hundreds of seconds; Figure 1C and 1F). The Kar9+Bim1+Bik1 and especially Kar9+Bim1 droplets became rapidly viscous; droplets fusing late in the reaction did not relax but formed chains of joint droplets (Figure 1C). Thus, the composition and aging properties of droplets widely affected their fluidity.

The ability of macromolecules to phase separate relies on the presence of a high valency of low affinity interactions between condensate components (reviewed in (Banani et al., 2017; Wheeler and Hyman, 2018)). Our data indicate that Kar9, Bim1, and Bik1 must indeed share multiple interaction interfaces with themselves and with each other. We therefore aimed next at dissecting some of these interfaces.

### Self-association of Kar9

The crystal structure of the folded N-terminal domain of Kar9 (Kar9N) from the budding yeast *Naumovozyma castellii* (NcKar9N) revealed three crystallographic dimers with total buried surface area per protomer of 1549, 1177, and 851 Å^2^, respectively (Kumar et al., 2021); the interfaces forming these dimers are referred to as α, β, and γ (Figure 2A). While the α and β interfaces are characterized by two symmetric contact points, the γ interface is established by a single one (Figure S1A-C). Notably, the residues forming the three interfaces (see Materials and Methods) are well conserved across Kar9 orthologues (Figure S2).

**Figure 2:**
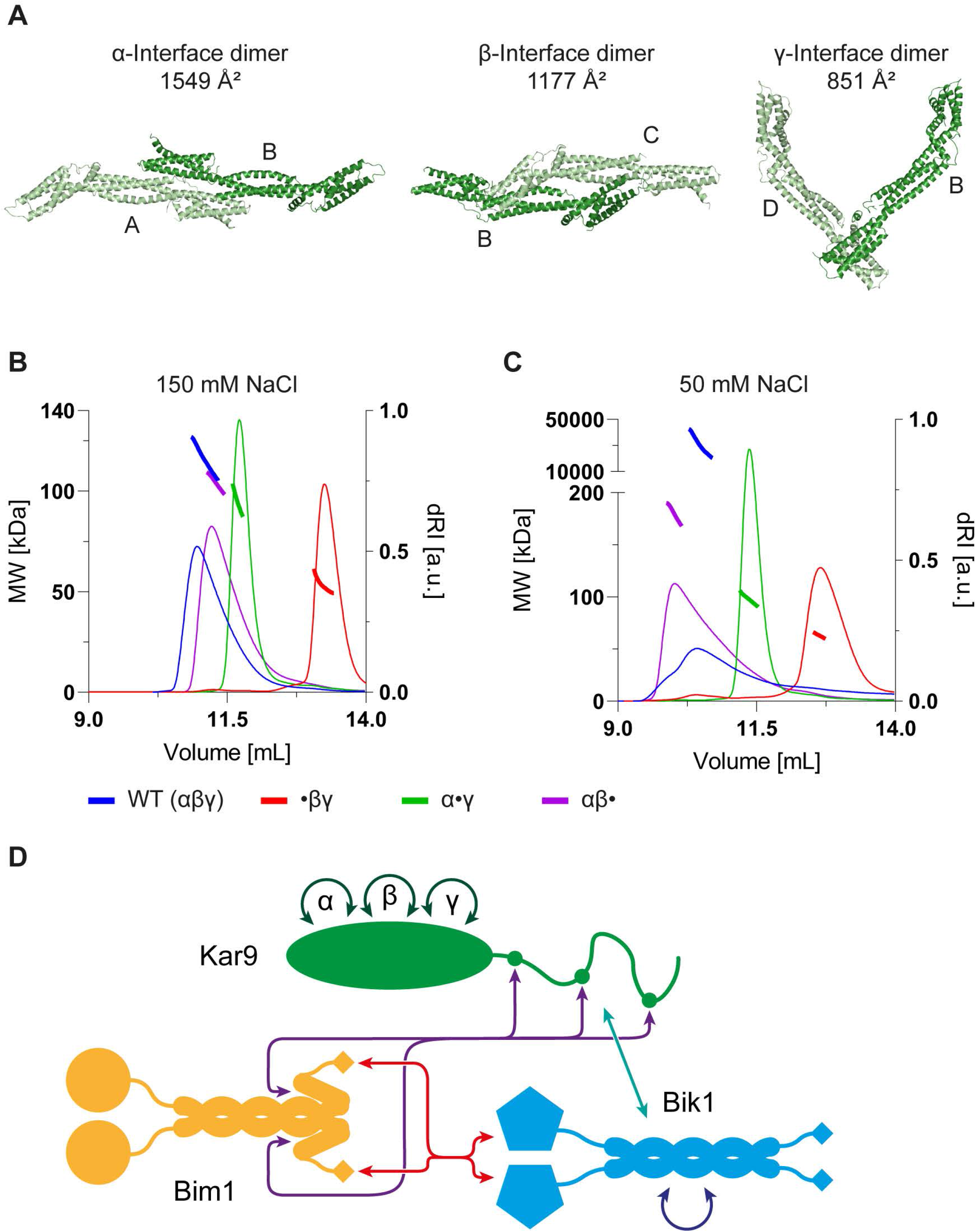
Characterization of NcKar9N-NcKar9N interactions. **A**, The three crystallographic dimers observed in the NcKar9N crystal (PDB ID 7AG9) in ribbon representation. The interfaces mediating the three dimers are referred to as α, β, and γ. The protomers in each crystallographic dimer are colored in light and dark green, respectively, and the buried surface area for each dimer is indicated. The four NcKar9N protomers that form the crystal are denoted A, B, C, and D. **B** and **C**, SEC-MALS experiments of His-tagged NcKar9N-αβγ (wild type; blue), NcKar9N-•βγ (α-interface mutant; red), NcKar9N-α•γ (β-interface mutant; green), and NcKar9N-αβ• (γ-interface mutant; magenta) in the presence of 150 mM (B) or 50 mM (C) NaCl. The Greek letters indicate the three intact interfaces; the dots indicate the mutated interfaces. The mutations are as follows: NcKar9N-•βγ, Phe288Ala/Phe344Ala;NcKar9N-α•γ, Arg233Ala/Ile237Ala;NcKar9N-αβ•, Tyr363Ala/Arg364Ala (see also Figure S1A-C). Protein solutions at a concentration of 200 μM were injected onto the SEC column. **D**, Schematic representation of the core Kar9-network module. Known protein-protein interactions between Kar9, Bim1, and Bik1 are indicated by double arrows.

To test whether any of the three interfaces could mediate NcKar9N-NcKar9N interactions in vitro, we performed a mutagenesis study with His-tagged NcKar9N. To this end, we mutated two conserved residues in each of the crystallographic dimer interfaces to alanine (Figures 2B and S1A-C). The integrity of the three mutants was confirmed by circular dichroism spectroscopy, which revealed spectra and cooperative thermal unfolding profiles similar to the wild type protein (Figure S1D and S1E).

The oligomerization state of the different NcKar9N variants was then assessed by size exclusion chromatography followed by multi-angle light scattering experiments. Consistent with our previous results (Kumar et al., 2021), analysis of wild type NcKar9N (200 μM protein injected onto the size exclusion chromatography column) at 150 mM sodium chloride yielded molecular masses consistent with the presence of dimers (118 kDa; calculated molecular mass of wild type His-NcKar9N monomer is 50 kDa; Figure 2B). At 50 mM salt, a broader elution profile shifted to a lower elution volume. The measured molecular mass increased to a value way above 200 kDa, suggesting the formation of non-stoichiometric oligomers (Figure 2C). In both salt conditions, mutating the α interface resolved these multimers to monomers (52 kDa). Whereas the β-interface mutation did not affect the dimer formed at 150 mM salt, it shifted the larger oligomers formed at 50 mM salt towards a molecular mass close to that of a dimer (97 kDa). The γ-interface mutant reached a mass of 180 kDa under both salt conditions, consistent with limiting the formation of oligomers up to tetramers. Together, these observations confirm that all three crystallographic homodimer interfaces mediate NcKar9N-NcKar9N interactions of different strengths, with the α interface mediating the most stable interaction.

As illustrated in Figure 2D, the C-terminal domain of Kar9 contains at least three binding sites for the C-terminal domain of the Bim1 dimer (Kumar et al., 2021; Manatschal et al., 2016; Zimniak et al., 2009). Furthermore, previous studies have indicated that Kar9 carries at least one binding site for the Bik1 dimer in its C-terminal domain (Moore et al., 2006), which is consistent with our finding that Kar9 phase separates together with Bik1 (Figure 1C). Since Bik1 also undergoes phase separation on its own (Figure 1C; (Ijavi et al., 2021)), it must contain at least three homodimerization interfaces. Additionally, the CAP-Gly domain of Bik1 was shown to interact with the C-terminal tail of Bim1 (Stangier et al., 2018). Thus, Kar9, Bim1, and Bik1 are indeed linked together by a dense network of interactions (Figure 2D).

### Role of Kar9 self-association for phase separation of the core Kar9-network components in vitro

We next reasoned that the network of multivalent interactions between Kar9, Bim1, and Bik1 (Figure 2E) probably drives its phase separation in vitro and if so that they should contribute cooperatively to this phenomenon. We thus rationalized that mutating interaction sites individually may have some effect on the ability of the network to undergo phase separation, but that accumulating such mutations should additively impair coalescence of network components. To test this idea, we inferred in *Saccharomyces cerevisiae* Kar9 the self-association sites identified in NcKar9N based on sequence conservation, and mutated them individually or in combination (see Materials and Methods and Figure S2). We then investigated whether these mutations affected the phase separation behavior of a 1:1:1 Kar9+Bim1+Bik1 mixture.

As shown in Figure 3A, interfering with any of the Kar9 self-association sites individually had some effect on the ability of the protein to promote phase separation, as shown by the fact that the corresponding phase diagrams deviated from that of the wild type protein at low protein concentration (1.0 μM) and at high sodium chloride concentrations (>300 mM). Among the three interface mutations, interfering with the γ interface had the strongest effect at high salt and protein concentrations (>300 mM and 10 μM, respectively), followed by mutation of the α interface. At low salt and protein concentrations (<200 mM and 1 μM, respectively), the α-interface mutant showed the strongest effect. The β-interface mutant behaved very similarly to wild type Kar9, except that it did not promote phase separation at lower protein concentration (1.0 μM). Mutating the three interfaces together enhanced these effects, although the protein was still able to promote phase separation of the protein mixture over a reduced but still broad range of concentrations and salinity (Figure 3A and 3B). Strikingly, for each of these variants, removal of Bik1 strongly reduced their ability to promote phase separation. When only Bim1 and Kar9 were present, the triple Kar9 interface mutant showed a much stronger coalescence defect compared to any of the single mutants and allowed to the formation of only few and small droplets (Figure 3A and 3C).

**Figure 3:**
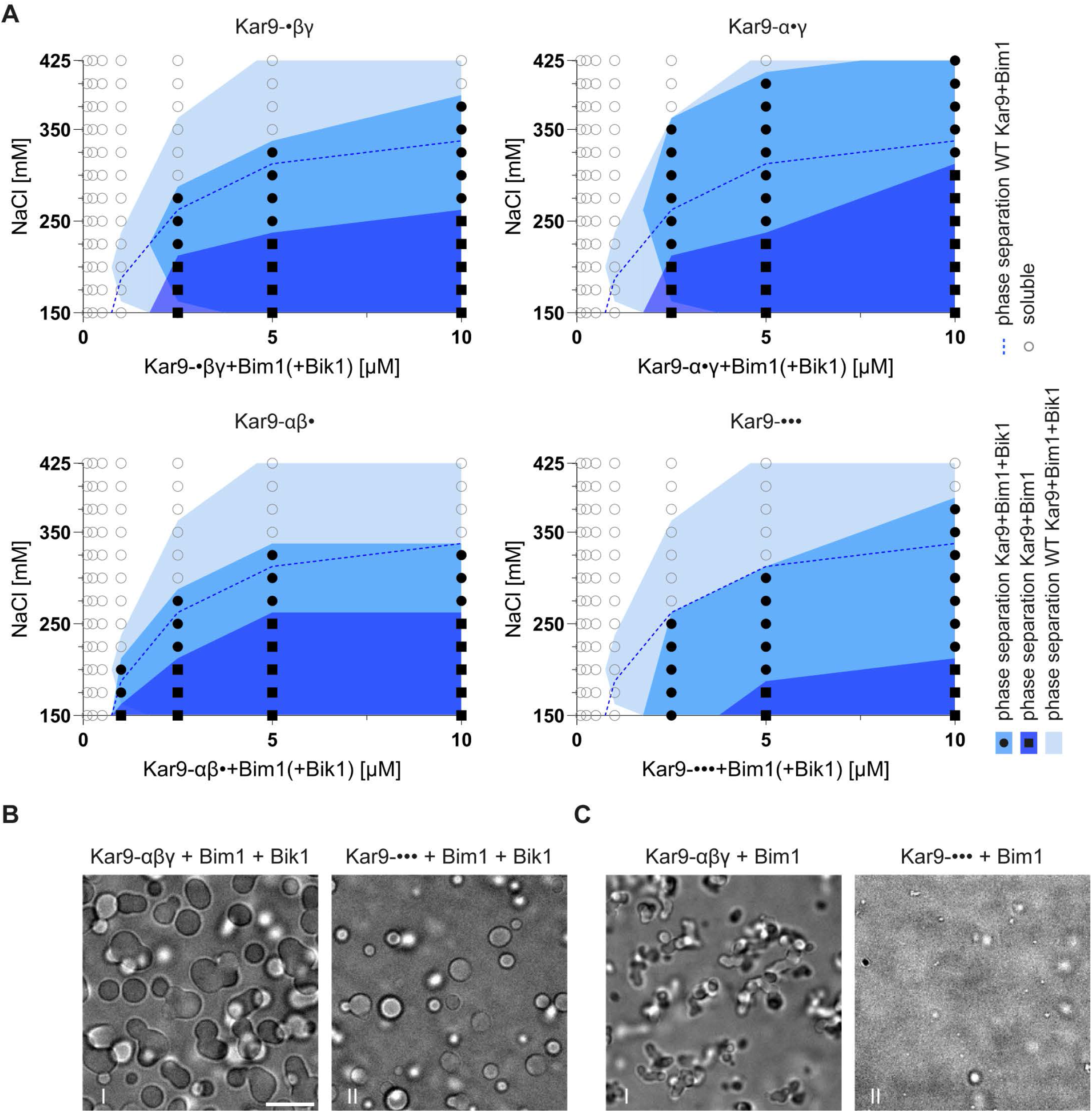
Impact of mutations in the core Kar9-network components on phase separation in vitro. **A**, Superimposed phase diagrams of different Kar9-interface mutants with Bim1 in the presence (intermediate blue) or absence (dark blue) of Bik1. For comparison, the phase diagrams of wild type Kar9+Bim1+Bik1 (light blue) and Kar9+Bim1 (dashed dark blue line) equimolar mixtures (Figure 1D) are also shown in all plots. **C** and **B**, Phase contrast microscopy images comparing phase separation of 10 μM wild type Bim1 with wild type Kar9 (I) or the triple homodimerization Kar9 interface mutant in the presence (B) or absence (C) of Bik1 (II) at 220 mM NaCl. Scale bar, 10 μm.

Thus, Bik1 and the three homotypic Kar9 interaction sites contribute each to the phase separation behavior of the core Kar9-network components in a cooperative manner. Since strong defects were observed only when combining Kar9 mutations and removing Bik1, we concluded that the different interactions between Kar9, Bim1, and Bik1 act redundantly in coalescence. Although we have certainly not identified all interfaces linking the three proteins together, our data indicate that simultaneously abrogating the identified ones is already sufficient to perturb coalescence in vitro.

### Role of Kar9 self-association for Kar9-network function in vivo

Next, we investigated whether the identified interfaces contribute to Kar9-network function in vivo. To this end, we constructed mutant strains where the endogenous *KAR9* gene was replaced with alleles abrogating each homotypic Kar9 interaction interface alone, in combinations of two, or all three together, in the presence or absence of Bik1. Since Kar9 function becomes essential for growth in cells lacking the dynein gene *DYN1* (Miller and Rose, 1998) each of the Kar9 alleles was combined with the *dyn1Δ* mutation to assay their functionality. Since beyond its role in the Kar9 network Bik1 is also essential for dynein function (Miller and Rose, 1998; Sheeman et al., 2003), we did not need to remove the *DYN1* gene in the *biklΔ* mutant cells to assay Kar9 functionality. As shown in Figures 4A and S3 mutating all three Kar9 interfaces and deleting Bik1 was lethal or nearly so over the entire range of temperatures tested (25-35°C), indicating that in these cells the Kar9 network was not functional. Remarkably, however, mutating any of the α, β, and γ interfaces or removing Bik1 individually did not cause much of a detectable phenotype at any of these temperatures, even when the interface mutations were combined with the *dynlΔ* mutation (Figures 4A and S3). Furthermore, any combination of two of these mutations failed to cause a much detectable phenotype, except eliminating the α interface and Bik1 together, which caused the cells to become sensitive to temperatures above 33°C. Thus, only some combinations of three mutations and removing the three Kar9 self-interaction interfaces and Bik1 together (the quadruple mutant) affected profoundly the growth and viability of the cells over a broad range of temperatures. We concluded that all three Kar9 self-interaction interfaces are functional in vivo, albeit redundantly with each other and with Bik1.

**Figure 4:**
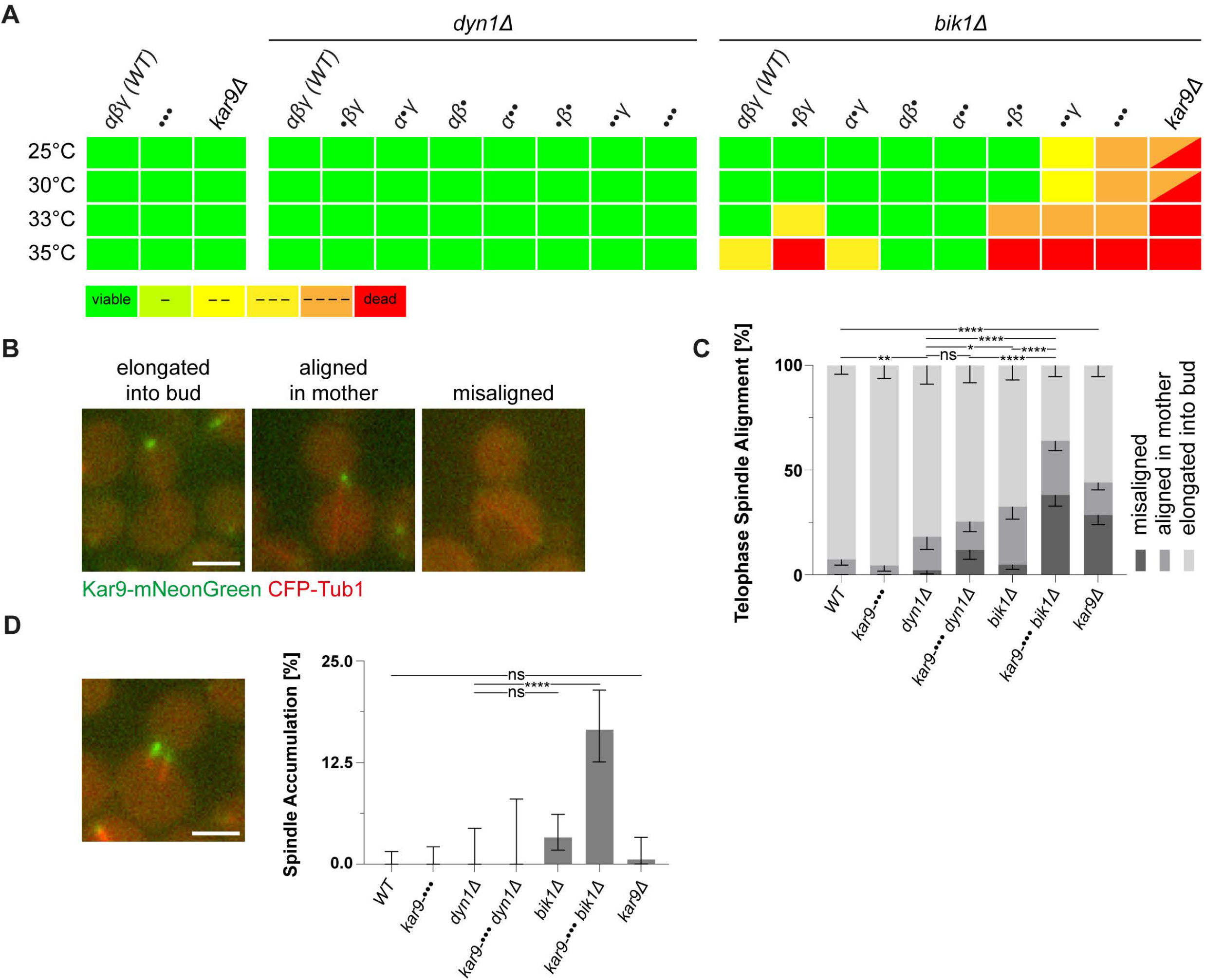
Role of Kar9-network multivalency for cell division. **A**, Summary of viability of Kar9 self-association interface mutants in combination with deletion of dynein or Bik1 at different temperatures as determined by spot assays (Figure S3). Viability is indicated by a color code from green (viable) to red (inviable). **B**, Micrographs showing anaphase spindle (CFP-Tub1, red) alignment of cells expressing Kar9 wild type or interface mutants tagged with mNeonGreen (green). Scale bar, 3 μm. **C**, Frequency of anaphase spindle alignment as shown in panel (B) (error bars are Wilson/Brown 95% confidence intervals). Significance tests compare the frequency of spindles elongated to the bud by two-proportion z test. Significance levels here and in figures 5–6: ns, p > 0.05; *p < 0.05; **p < 0.01; ***p < 0.001; ****p < 0.0001. **D**, Example image and quantification of frequency of pre-anaphase cells with more than one spindle (error bars are Wilson/Brown 95% confidence intervals). Significance levels determined by two-proportion z test. Scale bar, 3 μm.

To characterize how the progressive reduction of multivalency in the Kar9 network affected cell viability, we investigated the effects that cumulating mutations had on positioning the mitotic spindle (Figure 4B and 4C). While the *biklΔ* and the Kar9 triple-interface mutations had separately no or little effects on the outcome of mitosis at 25°C, their combination caused the accumulation of telophase cells with fully elongated spindles that remained in the mother cells and failed to deliver an SPB and the associated nucleus to the future daughter cell. Cells combining the Kar9 triple-interface mutant allele with the *dynlΔ* mutation showed an increased frequency of mispositioned telophase spindles but not nearly as much as the cells lacking the three Kar9 self-interfaces and Bik1 together, two thirds of which failed to properly segregate an SPB into the bud. Accordingly, up to 17% of these quadruple mutant cells contained more than one spindle during mitosis, a phenotype that is essentially never observed in wild type cells (Figure 4D).

Thus, the multivalency of interactions that drive network coalescence in vitro is also instrumental for the Kar9 network to guide properly the positioning of the mitotic spindle in vivo. Furthermore, like for phase separation in vitro, the functions of Bik1 and each of the homotypic Kar9 interfaces are largely redundant with each other, indicating that they act in concert in spindle positioning.

### Role of Kar9 self-association in Kar9 recruitment to microtubule tips in vivo

To understand how multivalency supports the function of the Kar9 network at the molecular level, we investigated whether it contributes to the localization of the Kar9 protein in vivo. At 25°C, cells expressing wild type or the mutated variants of Kar9 tagged with the fluorescent protein mNeonGreen revealed that erasing the γ interface had little effect on Kar9 localization but mutating the α or β interface reduced Kar9 levels at microtubule plus-ends by 40% and 20%, respectively (Figure 5A). Simultaneously erasing an increasing number of interfaces further decreased the levels of Kar9 at microtubule tips. Abrogating all three interfaces reduced these levels by 60%, and by 70% when Bik1 was removed as well. Strikingly, inactivating all three homodimerization interfaces of Kar9 also affected the ability of Kar9 to remain focused on the tip of one microtubule. In these cells, Kar9 was more frequently observed on both sides of the spindle (30% versus 10% in the wild type), a phenotype that is enhanced upon *BIK1* deletion (Figure 5B). Thus, the multivalency of interactions in the Kar9 network contributes to Kar9 recruitment to microtubule tips as well as to limiting the number of microtubules to which it is recruited.

**Figure 5:**
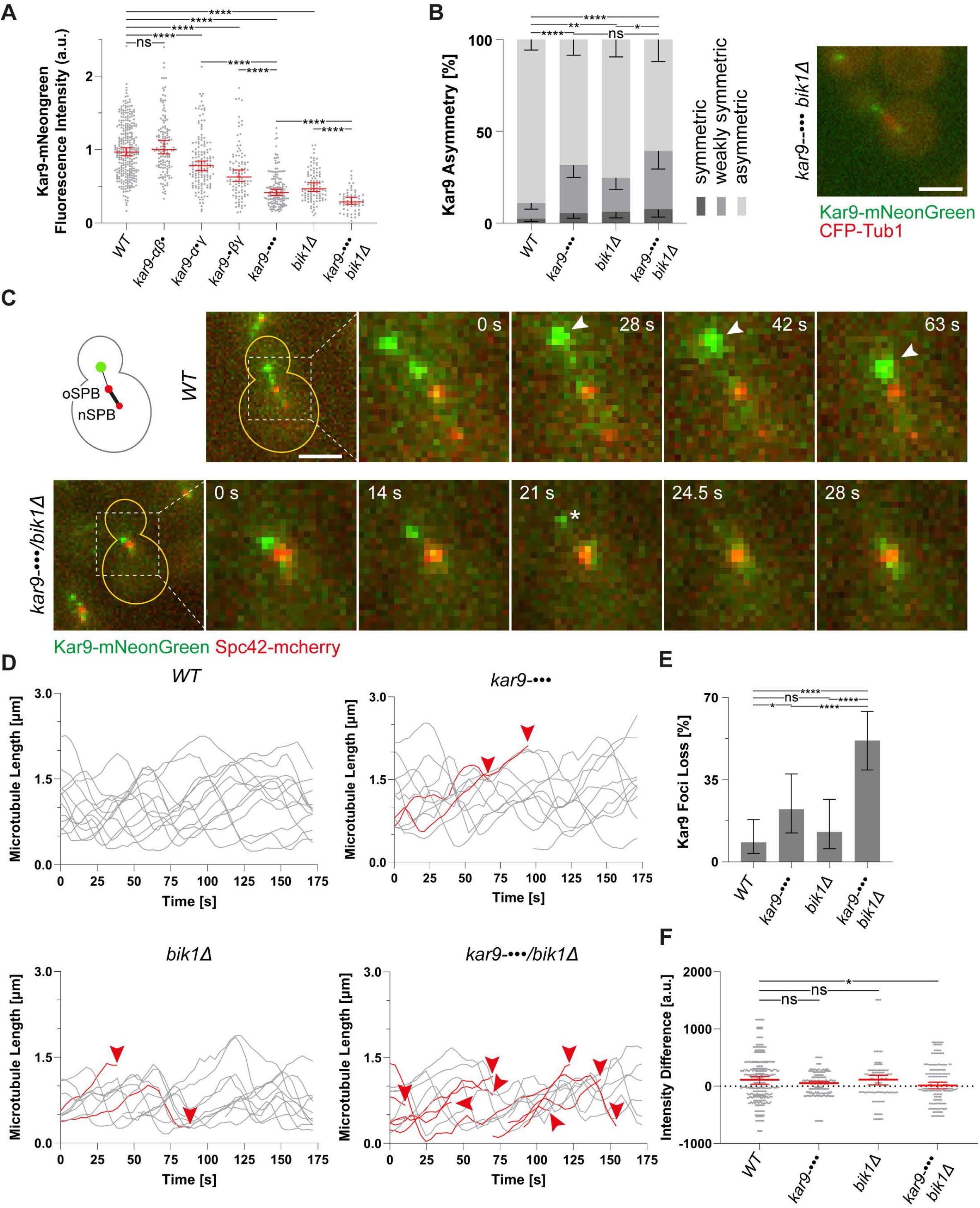
Dissection of Kar9 pathway defects caused by reduced Kar9-network multivalency. **A**, Kar9-mNeonGreen fluorescence intensity levels in Kar9 self-association interface mutants and Bik1 delete (bik1Δ; normalized median with 95% confidence intervals). Significance levels determined by Welch’s t-test. **B**, Frequency of asymmetric Kar9-mNeonGreen distribution between astral microtubule tips on both sides of metaphase spindles (error bars are Wilson/Brown 95% confidence interval). Significance levels determined by two-proportion z test. Example for weakly symmetric Kar9-•••/bik1Δ cell shown on the right. Scale bar, 3 μm. **C**, Micrographs showing Kar9-mNeonGreen tracking the tip of a shrinking astral microtubule at 25 and 37°C in wild type (arrowheads mark the period of shrinkage) and failure to do so in a Kar9-•••/bik1Δ cell at 25°C and Kar9-•••/dyn1Δ cell at 37°C (asterisk indicating loss of Kar9 focus). Scale bar, 3 μm. **D**, Astral microtubule length profiles for wild type, Kar9-•••, bik1Δ, and Kar9-•••/bik1Δ cells (moving average of distance between Kar9-mNeonGreen and Spc42-mcherry over three time frames). Profiles persisting for the whole length of the movie are displayed in grey, profiles that end due to loss of dot (indicated with arrowhead) in red. **E**, Frequency of cells losing Kar9-mNeonGreen focus on astral microtubule plus-end over 175 s (error bars are Wilson/Brown 95% confidence intervals). Significance levels determined by two-proportion z test. **F**, Kar9-mNeonGreen intensity change rate for shrinking astral microtubules, weighted by number of time points of shrinkage (median with 95% confidence interval, see methods section for details). Significance levels determined by Wilcoxon rank sum test.

### Role of Kar9-network multivalency for microtubule-tip tracking in vivo

The Kar9 network possesses the unique ability to track the tip of microtubules not only during their growth but also during shrinkage (Kusch et al., 2002; Liakopoulos et al., 2003). We thus wondered whether multivalency also contributes to this functionality, possibly by providing avidity of the Kar9-network components for growing and shrinking microtubule tips. To test this idea, we imaged the wild type and triple interface mutant forms of Kar9 tagged with mNeonGreen at high spatial and temporal resolution at 25°C. As previously shown (Liakopoulos et al., 2003), the dot formed by wild type Kar9 at the tip of mainly one microtubule per cell tracked this tip throughout the growth and shrinkage cycles of the microtubule (Figure 5C and 5D). Other Kar9 dots formed occasionally on microtubules but rapidly disappeared by either fusing with the main dot or by fading away in the cytoplasm.

This processive tracking of growing and shrinking microtubules by Kar9 was occasionally lost in cells where Kar9’s three homodimerization interfaces were simultaneously abrogated and in *biklΔ* mutant cells (Figure 5D-E). Bringing these mutations together boosted the phenotype: nearly 50% of the mutant cells lost their main Kar9 dot within a three-minute window, while less than 9% of wild type cells did so (Figure 5D-E). In nearly all cases, the dots reassembled next to the SPB, from where they subsequently tracked the next emerging microtubule plus-end. Dot disappearance generally happened at what seemed to be the end of a microtubule growth phase or shortly after shrinkage started. Consequently, specifically in the cells where the three interfaces were abrogated and Bik1 was not present the average intensity of the Kar9 dot did not increase during microtubule shrinkage, as it did in the wild type cells during that time (Figure 5F). Again, this behavior was not observed at all or not nearly as clearly in cells carrying either the *biklΔ* or the Kar9 triple interface mutation alone.

Thus, these results show that Bik1 and the three homodimerization interfaces of Kar9 act in concert in the focused recruitment of Kar9 to virtually a single selected microtubule tip in vivo. They also establish that within these foci, a substantial fraction of Kar9 is recruited not through microtubule-bound Bim1, but through interaction with itself and with Bik1. Notably, the multivalent interactions in the Kar9 network provide a molecular basis to understand the network’s ability to keep tracking a microtubule end not only during growth but also as it shrinks.

### The Kar9 network forms a non-stoichiometric structure at microtubule tips

Since the function of Kar9 in vivo (Figures 4 and 5) correlates very well with its ability to promote the formation of liquid-like droplets together with Bim1 and Bik1 in vitro (Figure 1–3), we hypothesized that the three proteins might function together by forming a condensate at the tip of their target microtubule in vivo. However, in vivo the Kar9 dots at microtubule tips are smaller than the diffraction-limited resolution of light microscopy, preventing to visualize their morphology and directly assaying their material properties. Thus, we investigated whether they showed other hallmark properties of biomolecular condensates. One such hallmark property consists of a condensate’s ability to vary in stoichiometry and size as the amount of available material changes (reviewed in (Banani et al., 2017)). Thus, we asked whether Kar9 dots showed such behavior in vivo, using Dam1 as a reference, which forms a stoichiometric complex at the tip of kinetochore microtubules (Jenni and Harrison, 2018; Joglekar et al., 2006).

As shown in Figures 6A, 6B and S3, Dam1 tagged with mNeonGreen formed dots of constant fluorescence intensity, irrespective of cell cycle stage or Dam1 expression levels. In contrast, the dots formed by Kar9-mNeonGreen widely varied in intensity as cells progressed from G1 to metaphase, upon insertion of a second copy of the gene in the genome or when over-expressed (Figures 6A-C). Moreover, abrogating all three homotypic interfaces of Kar9 reduced about four-fold the intensity of the dots that it formed upon overexpression (Figure 6B-C). Thus, the Kar9 network does not assemble as a stoichiometric structure at the tip of a microtubule, but condenses into a cohesive structure. Dam1 is present in 17 copies per kinetochore microtubule (Jenni and Harrison, 2018). Using its intensity as a reference, we estimate that in metaphase cells there are at least 70 Kar9 molecules on average at the tip of a single microtubule (see Materials and Methods).

**Figure 6:**
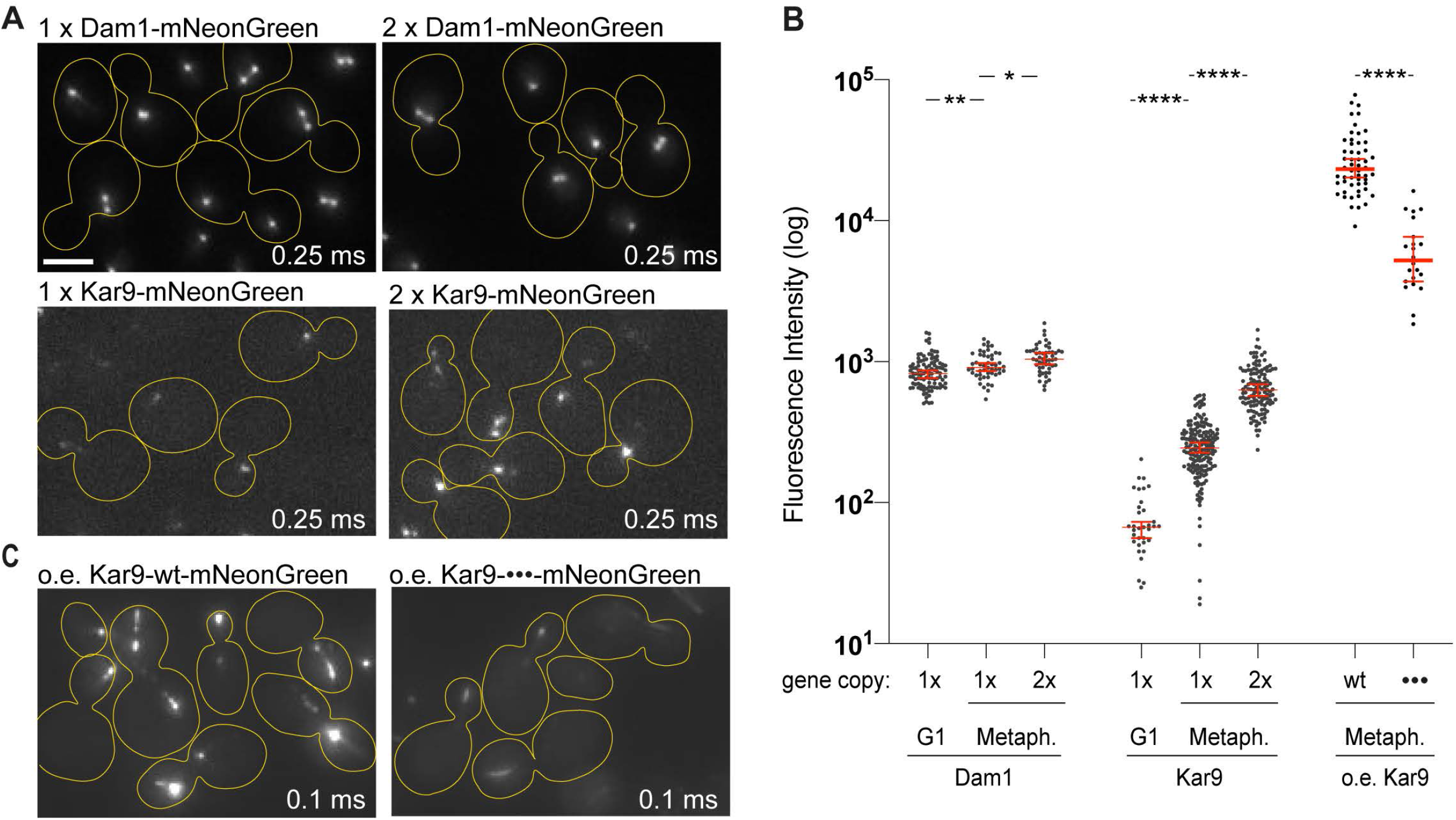
Kar9 stoichiometry at microtubule tips. **A**, Micrographs showing Dam1-mNeonGreen or Kar9-mNeonGreen localization in haploid cells bearing one (1x) or two (2x) fluorescently tagged gene copies (exposure time displayed in ms; scale bar 3 μm). **B**, Fluorescence intensity levels of Dam1-mNeonGreen and Kar9-mNeonGreen during G1- and metaphase, of the cells described in panel (A) (median with 95% confidence interval; significance levels determined by Welch’s t-test). **C**, Micrographs showing the localization of Kar9-mNeonGreen overexpressed from the galactose promoter in haploid cells (exposure time displayed in ms).

### Kar9 shows liquid-like behaviors on microtubules in vivo

We observed that Kar9 dots frequently merged with each other in cells containing several ones, even when Kar9 was expressed at endogenous levels (Figure 7A). Subsequently, these merged dots remained stably together for many time frames, suggesting that they had truly fused with each other into a single entity. These observations suggest that the Kar9 network functions as a liquid-like condensate.

**Figure 7:**
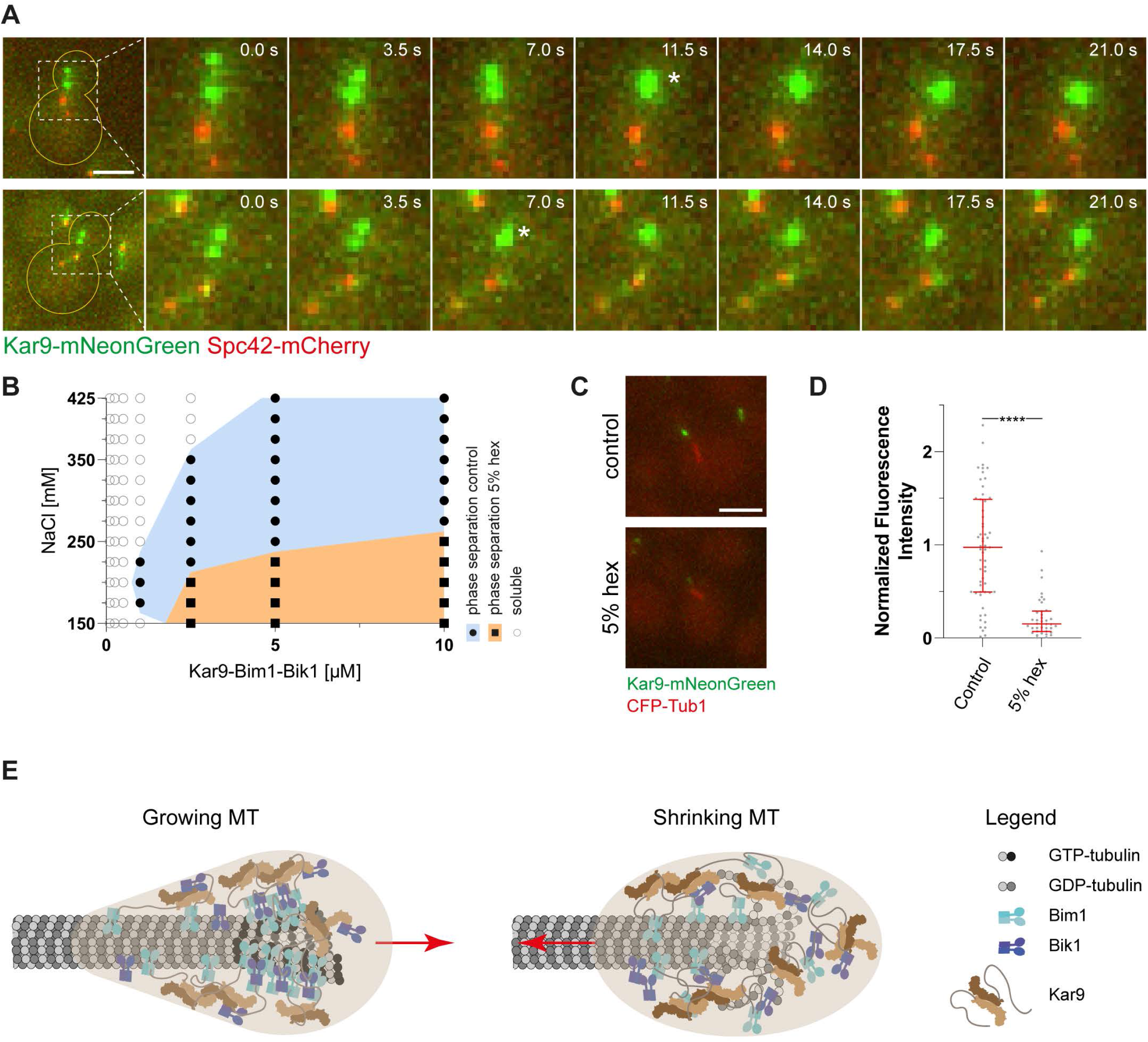
Kar9 liquid-like behavior in vivo. **A**, Micrographs showing fusion events between Kar9-mNeonGreen foci. Scale bar, 3 μm. **B**, Superimposed phase diagrams comparing the phase separation of 1:1:1 mixtures of Kar9+Bim1+Bik1 in presence or absence of 5% 1,6-hexanediol (hex). **C**, Micrographs of a control cell and cell treated with 5% 1,6-hexanediol immediately before acquisition. Scale bar, 3 μm. **D**, Kar9-mNeonGreen fluorescence intensity levels in wild type cells treated with 5% 1,6-hexanediol versus untreated control (normalized median with 95% confidence intervals). Significance levels determined by Welch’s t-test. **E**, Proposed model for how the Kar9 body tracks growing (left) and shrinking (right) microtubules.

To further test the molecular mechanisms driving the assembly of Kar9 dots, we characterized their sensitivity to hexanediol. This alcohol has been reported to resolubilize certain liquid-liquid phase separated proteins both in vitro and in vivo, while leaving other stoichiometric complexes and aggregates mostly intact (Kroschwald and Alberti, 2017). Indeed, phase separation of Kar9+Bim1+Bik1 droplets was severely affected in vitro by the addition of 5% hexanediol, especially at medium sodium chloride concentrations (>250 mM; Figure 7B). Similarly, the intensity of Kar9 dots in vivo was reduced about eight-fold when imaged immediately after suspending the cells in medium containing the same amount of hexanediol (Figure 7C-D). Imaging of the mitotic spindle in the same cells, using CFP-tubulin as a reporter, indicated that the spindle remained stable under this treatment. These results support the notion that the high multivalence of low affinity interactions in the Kar9 network promotes the formation of a non-stoichiometric, extensible assembly with liquid-like properties at the plus-end of target microtubules in living cells.

## Discussion

A wide range of biomolecules use low affinity, multivalent interactions to undergo liquid-liquid phase separation and generate membrane-less organelles, including the nucleolus, centrosomes, and processing bodies (reviewed in (Wheeler and Hyman, 2018)). The liquid-like properties of these bodies are thought to provide a local environment where specialized molecules are enriched and others are excluded, thus either enhancing or inhibiting the activity of specific biosynthetic pathways (reviewed in (Banani et al., 2017; O’Flynn and Mittag, 2021)). However, it remains generally difficult to demonstrate firmly whether the condensate nature of bodies is determinant for their function - the mere fact that a set of proteins undergoes phase separation in vitro is no proof that they also do so in vivo. Moreover, due to the redundancy of the underlying interactions it is intrinsically challenging to perturb the condensation of a given system in a controlled manner in vivo to determine whether condensation contributes to function, and if so to which ones, and how.

Here, we show that a dense web of redundant interactions links the core components of the Kar9-network together to assemble a structure, which we name the “Kar9 body”, at the plus-end of essentially a single cytoplasmic microtubule in metaphase cells of budding yeast. Our data suggest that this body is a liquid-like condensate, the dynamic and microtubule-binding properties of which allows it to persist and track both the growing and shrinking plus-end of a selected microtubule in vivo. We postulate that the material properties of the Kar9 body supports its function as a “mechanical coupling device” that supports the transfer of forces between the actin and the microtubule cytoskeleton to position the mitotic spindle for proper cell division.

Several lines of evidence support our hypothesis that the Kar9 body behaves as a liquid-like structure. Firstly, the phase separation of the core components of the Kar9 network together in vitro relies on multiple medium and low affinity interactions between them. The interfaces that we have identified in Kar9 so far, i.e., Kar9-Kar9 ((Kumar et al., 2021) and this study) and Kar9-Bim1 (Kumar et al., 2017; Manatschal et al., 2016) are all conserved across Kar9 homologs and, therefore, all functionally relevant. Secondly, the multivalent interactions within the Kar9 network are instrumental for its proper function. In this context, we found that their redundancy in vivo perfectly parallels their cooperativity for network phase separation in vitro. Thirdly, supporting the idea that the Kar9 body is a liquid-like structure in vivo, it is a non-stoichiometric assembly, which is able to undergo fission and fusion. These data support two key notions namely that the core components of the Kar9 network undergo condensation in vivo, and that this property is instrumental for the Kar9 network to fulfil its cellular functions.

An immediate prediction emerging from this view is that the phenotypic and molecular consequences of decreasing multivalency in the system must reveal which body’s functionalities are specifically dependent on phase separation. Our data provide compelling support for condensation mediating at least two key features of Kar9 network function. Firstly, our in vivo results indicate that condensation underlies the recruitment and concentration of Kar9 to a selected microtubule tip in vivo. Kar9 has a high affinity for Bim1, and Bim1 for the plus-end of growing microtubules (Manatschal et al., 2016; Maurer et al., 2012). Thus, the recruitment of Kar9 to microtubule tips via Bim1 must locally enhance its concentration and may nucleate condensation, leading to the recruitment of more Kar9 and of its partners. Oswald ripening of this condensate, i.e., the tendency of the biggest condensate to collect all the available material, could explain why cells form one and only one Kar9 body in their cytoplasm (Liakopoulos et al., 2003). Thus, condensation offers a determining mechanism for microtubule specialization.

Secondly, multivalency underlies the remarkable capacity of the Kar9 body to track the plus-end of shrinking microtubules, contributing thereby to the cohesion and persistence of Kar9 to a single microtubule over time in vivo. As condensation proceeds, accruing Kar9 levels should bring an excess of Bim1 molecules with itself, allowing Bim1 binding to low affinity sites on the microtubule lattice (Geyer et al., 2015). Supporting this notion, we found that overexpression of Kar9 tends to extend the Kar9 body along the microtubule lattice. We suggest that such “wetting” of the lattice supports both the persistence of the Kar9 body on shrinking microtubule plus-ends (Figure 7E), as well as its cohesion to the dynamic microtubule tip necessary to support the transmission of pulling forces generated either by microtubule shrinkage or by the directional movement of myosin motors along actin cables. Thereby, condensation should facilitate the function of the Kar9 body in mechanically linking a microtubule plus-end with an actin cable to mediate nuclear orientation and positioning.

The emerging effects of Kar9 network condensation underlie the broader role of multivalency in enabling the proper segregation of one and only one spindle pole to the bud. The hypothesis of a liquid-like Kar9 body is especially attractive because it helps explain many cellular observations that have remained difficult to rationalize so far, such as Kar9’s unique localization pattern and its remarkable role in “gluing” together two highly dynamic structures, namely microtubule tips and actin cables. Similar mechanisms may as well explain how other patterning +TIPs, such as APC and MACF, target, persist, and specialize the dynamics and functions of specific subclasses of microtubules in other cell types. However, beyond their relevance for the organization of microtubule networks and the specialization of individual microtubules, our findings also raise the question of how small can functional biocondensates be inside cells. Indeed, the Kar9 body described here is much smaller than typical membrane-less organelles. We have estimated that in metaphase cells it is formed of perhaps as few as 70 Kar9 molecules; probably similar numbers apply for Bim1 and Bik1. Thus, our study raises the question of how many molecules are needed to assemble a “nano-fluid”, how their organization compares to that of bigger droplets, and how it supports their biological functions. Further studies will be necessary to investigate whether such “nano-fluids” are widespread in living matter, the spectrum of functions they may fulfill, as well as how their size contributes to their properties.

While we were assembling our manuscript, we learned about the work of three additional groups, which describes the phase separation of +TIP systems from other organisms. Miesch et al. found in vitro and in cellulo that the two human +TIPs EB3 (related to Bim1) and CLIP-170 (Bik1) undergo phase separation at growing microtubule plus-ends, and by doing so concentrate tubulin to this location. As a result, microtubule dynamics was strongly affected. Song et al. demonstrate that EB1 (Bim1) phase separates on its own via an intrinsically disordered region, that the patterning +TIP TIP150 potentiates this process at kinetochores, and that this condensation process plays an important role in spindle assembly and function in human cells. Maan et al. in turn investigated in vitro the role of the multivalent interactions between the *Saccharomyces pombe* +TIPs Mal3 (Bim1), Tea2 (kinesin), and Tip1 (Bik1). The authors found that the intrinsically disordered regions of Mal3 drive the phase separation and thus the accumulation of Tip1 and Tea2 at growing microtubule tips. In combination with our study, this extensive amount of work converges into a compelling picture in which liquidliquid phase separation provides a general mechanistic framework to understand how +TIPs function at microtubule ends. It also indicates that this mechanism may come in diverse flavors. While phase separation of ubiquitous +TIPs might be universal at the tip of growing microtubules, patterning +TIPs, such as Kar9 and probably TIP150, are required for condensation to extend tracking to shrinking microtubules, to support microtubule specialization, and possibly to transmit mechanical forces. Studying further the condensate properties of +TIP bodies may thus change our understanding of how +TIPs accumulate to selected microtubule plus-ends, functionally specialize them, control their dynamics, and regulate their interactions with different cellular structures.

## Acknowledgements

We thank Madhurima Choudhury, Mathias Bayer and Toni Vukovic for their help with the experiments, Marcel Stangier for helping with protein production and ScopeM for their support and assistance with microscopy. We thank previous and current members of the Steinmetz, Dufresne and Barral laboratories for extensive and critical discussions throughout the project. This work was supported by a Marie Curie COFUND fellowship (to AK) and by grants from the Swiss National Science Foundation (31003A-105904 to YB and 31003A_166608 and 310030_192566 to MOS; Sinergia CRSII5_189940 to YB, MOS and ED) and from SystemsX.ch (RTD Grant #2012/192 TubeX, to MOS, JS and YB).

## Author contributions

S.M.M., A.-M.F. and A.K. designed and performed the research, and analyzed the data. M.I. characterized the material properties of the different droplets in vitro (Fig. 1F). J.S. designed and supervised the quantitative analysis of Kar9 levels on shrinking microtubules (Figure 5F). R.B. designed and performed the analysis of Kar9 overexpression with A.-M. F. E.D. contributed to designing the research and supervised the work of M.I. Y.B. and M.O.S. designed and supervised the research, analyzed the data, and wrote the manuscript with input from all authors.

## Materials and Methods

### Cloning and protein preparation

#### Plasmid preparation

Expression vectors were produced either by homologous recombination with PCR-amplified or synthetic genes and amplified backbones or site directed mutagenesis (QuikChange, Agilent). The wild type (Kumar et al., 2021) and mutant N-terminally hexa-histidine tagged *N. castelli* Kar9 N-terminal domain (NcKar9N, residues 1-410) were cloned into the pET-based expression vector PSPCm2 (Olieric et al, 2010), respectively. The interface mutations in *N. castellii* Kar9 (UniProt ID G0VE12) are as follows: α, Phe288Ala/Phe344Ala; β, Arg233Ala/Ile237Ala; γ, Tyr363Ala/Arg364Ala. Full-length wild type (Manatschal et al., 2016) and mutant N-terminally hexa-histidine tagged *S. cerevisiae* Kar9 (UniProt ID P32526) was cloned into the Acembl vector pACE (Bieniossek, 2009). The interface mutations in *S. cerevisiae* Kar9 are as follows: α, Phe292Ala/Leu347Ala; β, Arg237Ala/Asn241Ala; γ, Tyr366Ala/Arg367Ala. Full-length *S. cerevisiae* wild type and N-terminally hexa-histidine tagged *S. cerevisiae* Bim1 (Manatschal et al., 2016) (UniProt ID P40013), and *S. cerevisiae* Bik1 (UniProt ID P11709) were cloned into pET3d (Invitrogen) and PSTCm1 (Olieric et al, 2016), respectively.

#### Protein Expression and Purification

Sequence verified plasmids were transformed into BL21(DE3) *E. coli* cells for protein expression. To produce the proteins, liquid cultures were shaken in LB media containing the appropriate antibiotic until an OD_600_ of 0.6 was reached. Expression was induced by the addition of 0.75 mM isopropyl-β-D-thiogalactoside (IPTG). Induced cells were further incubated overnight at 20 °C. Cells were lysed using the sonication method or high-pressure homogenizers (Avestin Emulsiflex-C3 High Pressure Homogenizer or Microfluidics Microfluidizer) in 20 mM Tris-HCl, (pH 7.5), supplemented with 800 mM (full length Kar9 alone, Manatschal 2016) or 500 mM (all other proteins) NaCl. Cell debris was removed by centrifugation. Cleared cell lysates were filtered using a 0.45 μm filter. Affinity purification of His-tagged proteins was carried out at 4°C by immobilized metal affinity chromatography on Ni^2+^-Sepharose columns (Cytiva) according to the manufacturer’s instructions. For NcKar9N wild type and full length Kar9 (purified without Bim1), the hexahistidine tag was enzymatically cleaved with PreScission and TEV protease, respectively. All protein samples were further applied on a pre-equilibrated Superdex-75 or −200 size-exclusion chromatography column (Cytiva). For the production of different Kar9-Bim1 complex variants, wild type or mutant hexahistidine tagged Kar9 and untagged Bim1 were separately expressed in bacteria and mixed before lysis in 500 mM NaCl. The purity of recombinant proteins was confirmed by Coomassie-stained 12% SDS-PAGE, and the identities of the proteins were assessed by mass spectral analyses.

### In vitro phase separation assays

#### Pelleting assay

Protein stocks were mixed and diluted in 1.5 mL test tubes with 20 mM Tris-HCl, pH 7.5, supplemented with 500 mM NaCl to reach a protein concentration of 40 μM each in 5 μL total volume at 500 mM NaCl (722 mM for Kar9 and Kar9 + Bik1). Next, 15 μL of low (100 mM or 26 mM for Kar9 and Kar9 + Bik1) or high salt (500 mM or 426 mM for Kar9 and Kar9 + Bik1) Tris buffer was added to dilute the samples 4:1 and to reach a final concentration of 10 μM in 200 or 500 mM NaCl, respectively. The mixtures were then incubated in a thermomixer for 30 min at 25°C while gently shaking. After 10 min spinning at 16’900 g and 25°C, the supernatant was transferred into a separate tube. The invisible pellets were resuspended in 20 μL of the Tris buffer (containing either 200 or 500 mM NaCl). Samples were analyzed by Coomassie stained 10% SDS-PAGE.

#### Static light-microscopy assay for the assembly of phase diagrams

A dilution series of either 5X (wt proteins) or 4X (mutant proteins and Kar9+Bik1, due to concentration limitations) protein mixtures as above was prepared at high salt (usually 500 mM, except experiments with Kar9 purified alone 718 mM) to reach a broad range of protein concentrations (usually 0.1 – 10 μM final concentrations after dilution, thus either 0.5 – 50 or 0.4 – 40 μM before dilution). The reservoir wells of an MRC2 crystallization plate (Swissci) were then filled with buffers matching the final salt concentration of 150 – 425 mM NaCl. Next, the protein stock solutions were dispensed into the MRC2 plate’s drop wells and diluted 5:1 or 4:1 (depending on the protein stock concentration) in a total volume of 1 μL using a Mosquito pipetting robot (sptlabtech) with dilution buffers provided in a different plate to reach salt concentrations matching the reservoir wells. After 30 min incubation, all wells were imaged by phase contrast microscopy. The phase separation state of each well is represented by a symbol in the phase diagrams shown in Figures 1 and 3. An approximate binodal line was then drawn between phase separated and soluble conditions (through the middle points at each tested protein concentration).

#### Time-resolved microscopy assay

Either 5X or 4X protein mixtures were prepared and then diluted to 10 μM and 220 mM NaCl in a Nunc^®^ Lab-Tek^®^ II chambered 8-well coverglass. To verify enrichment of His-tagged protein in droplets, Atto-488-NTA was used at 1 μg/mL concentration. Samples were imaged on a DeltaVision personalDV microscope (Cytiva) at 60x or 100x magnification using transmission and FITC channels in a single plane close to the coverglass surface.

#### Droplet fusion analysis

In the inset of the figure S4, a typical fusion event of Bik1 is shown. To measure the fusion time τ, we assume that a uni-axially deformed droplet relaxes to its final spherical state.

Parameter 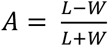, was used to calculate the fusion time τ.

L is the major axis of the deformed droplet and W is the minor axis. Fitting an exponential curve to the time dependent parameter A, figure (a), would give the fusion time τ.

τ has relationship with viscosity and surface tension via Navier-Stock’s equation:

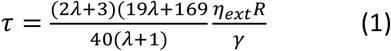

where *λ* = *η_int_*/*η_ext_* is the ratio of internal and the external viscosities, R is the final radius of the droplet in spherical state, and γ is the surface tension.

In the case that *λ* ≪ 1 (*η_int_* ≪ *η_ext_*), one can simplify the equation to:

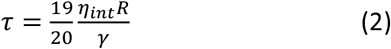

which is valid where the viscosity of the protein condensed phase (*η_int_*) is much bigger than the viscosity of the protein dilute phase (*η_ext_*). For instance, viscosity of the Bik1 droplet phase has been reported previously as *η_int_* ≈ 18.2 (*Pa* · *s*) which is four orders of magnitude higher than the Bik1 dilute phase (Ijavi et al., 2021).

In Figure 1F, 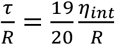(Eq. 2) is the slope measured from fitting the data linearly. The value of 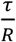 for Bik1 is measured as 2.25 ± 0.37 (*s/m*) which is consistent with the previously reported values of viscosity and surface tension for Bik1, 18.2 (*Pa · s*) and 7 (*μN/m*), respectively (Ijavi et al., 2021).

However, higher values of 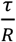 for Bik1+Kar9+Bim1 and Kar9+Bim 1, 1 1.34 ± 3.46 and 123.92 (*s/m*), suggests much slower fusion for these systems. Unfortunately, due to infrequency of fusion events for Kar9+Bim1 droplets, we could only resolve one event. Therefore, there is no meaningful error for this event reported (Jeon et al., 2018; Leal, 2007).

### Biophysical characterization

Size exclusion chromatography followed by multi-angle light scattering (SEC-MALS) experiments were performed in 20 mM TrisHCl, pH 7.5, supplemented with 150 or 50 mM NaCl, 1 mM DTT using a S-200 10/300 analytical size exclusion chromatography column (GE Healthcare) connected in-line to mini-DAWN TREOS light scattering and Optilab T-rEX refractive index detectors (Wyatt Technology). 30 μl of His-NcKar9N protein variants at a concentration of 200 μM were injected for each run onto the SEC column.

Circular dichroism (CD) spectra of His-NcKar9N protein samples were recorded at 5 °C and at a protein concentration of 0.2 mg/ml in 20 mM TrisHCl, pH 7.5, supplemented with 500 mM NaCl and 1 mM DTT using a Chirascan spectropolarimeter (Applied Photophysics) and a cuvette of 0.1 cm path length. Thermal unfolding profiles between 5 and 80 °C were recorded by increasing the temperature at a ramping rate of 1 °C/min monitoring the CD signal at 222 nm. Midpoints of thermal unfolding profiles were determined using the Global3 program (Applied Photophysics).

### Structural analysis of crystallographic NcKar9N homodimer interfaces

The three crystallographic NcKar9N homodimer interfaces were analyzed using the PDBePISA server (Krissinel and Henrick, 2007) using PDB ID 7AG9. The following residues form the three different interfaces: α, F181, Q184, E185, F188, F288, E292, E299, and K303, and Y318’, E322’, K329’, R343’, F344’, Q347’, K350’, and K351’; β, L107, E111, D123, F124, L127, and I131, and K179’, R233’, K234’, L236’, I237’, and R240’; γ, N360, Y363, R364, E367, R390, and L398, and N211’, N212’, Y363’, R364’, and L377’. The prime discriminates the second protomer of a respective crystallographic dimer. Figures were prepared using the software PyMOL (The PyMOL Molecular Graphics System, Schrodinger, LLC).

### Yeast strains and cloning

Yeast strains used in this study were created by transformation of PCR amplified fragments that integrated into the genome by homologous recombination (Kumar et al., 2021) or crossing, sporulation and tetrad dissection. Transformants were verified by PCR and sequencing, spores by replication onto relevant selection plates. Summary of all yeast strains in the key resources table.

### Yeast viability assay

Haploid yeast strains of the given genotypes were spotted in 10x serial dilutions, starting from OD_600_ 0.05, by spotting 4μl from the individual dilutions on YPD plates and incubating them at the indicated temperatures for 2-3 days.

### Yeast life-cell time-lapse fluorescent microscopy

For life-cell time-lapse fluorescent microscopy, yeast strains were exponentially grown in synthetic medium lacking tryptophan, harvested by centrifugation at 600 g for 1.5 min, and placed on glass slides for imaging. For the phenotypes telophase spindle alignment (Figure 4C), spindle accumulation (Figure 4D), Kar9 fluorescence intensity (Figure 5A), and Kar9 asymmetry (Figure 5B), cells were imaged on a Personal DeltaVision microscope (Cytiva) in CFP and FITC channels (*kar9Δ* only CFP) over 10 iterations separated by 13 sec in 17 z-slices separated by 0.3 μm. For microtubule plus-end tracking (Figure 5C-F), cells were imaged on a Visitron Spinning Disk microscope used in wide field mode with simultaneous acquisition in mCherry and GFP channels over 50 iterations separated by 3.5 sec in 17 z-slices separated by 0.3 μm.

For hexanediol treatment, cells were grown on YPD agar plates at low density and then resuspended in synthetic medium lacking tryptophan supplemented with 10 μg/mL digitonin, with or without 5% (w/v) 1,6-hexanediol. Imaging was performed as described above with a DeltaVision Personal microscope.

### Microscopy data analysis

For telophase spindle alignment, spindle accumulation, and Kar9 asymmetry, the seven most in-focus planes of the z-stack were sum projected and the phenotypes were manually classified at the second time point. The other time points were used to assign cells to the analyzed cell cycle stage.

For Kar9-mNeonGreen fluorescence intensity quantification, the complete z-stacks of time points 2-10 were maximum projected and the background (slide) subtracted maximum intensity for each cell was measured and averaged over the nine time points.

For microtubule plus-end tracking analysis, the complete z-stacks of all 50 time points were maximum projected. Kar9 focus loss (Figure 5E) was classified manually for these movies. Further, for each Kar9 focus the associated SPB was determined by looking at the trajectory of Kar9 fluorescence. The 2D distance between the Kar9-mNeonGreen focus and the relevant Spc42-mCherry focus (microtubule length) as well as the background subtracted (cell background) integrated fluorescence intensity was measured for each time point. Smoothed (moving average over 3 time points) microtubule length trajectories are shown in (Figure 5D).

Periods of microtubule growth and shrinkage were annotated automatically on non-smoothed trajectories. For each Kar9 focus, the procedure involved: (i) applying a Savitzky-Golay finite impulse response smoothing filter (polynomial order 1, frame length 5) to the microtubule length trajectory; (ii) identifying change points in the filtered data (max. 5 change points per trajectory, comparison to least squares linear fit, minimal distance between change points of 5 frames); (iii) detecting minima / maxima in the unfiltered trajectories closest to the identified change points (minimum required peak prominence of 25% of the difference between maximal and minimal microtubule length); and (iv) annotating periods as growth (shrinkage) when slopes of linear regressions on the partial trajectories between maxima / minima (or change points) were significantly positive (negative) based on t-tests on coefficients of linear regressions, with α=0.05. To estimate initial microtubule length or intensity as well as rate of microtubule length and fluorescence change for individual microtubule growth or shrinkage phases, we used ordinary linear regression on the respective partial trajectories. Reported values Figure 5E are the corresponding regression coefficients for fluorescence change during shrinkage, weighted by number of time points of the shrinkage event. Wilcoxon ranks sum test was used to calculate significance levels between medians. Automatic classification and regression were implemented in Matlab R2019b.

For stoichiometry analysis the z-stacks were maximum projected and Kar9 and Dam1 intensity (raw integrated density) was measured after subtracting the cell background. For estimating the brilliance of one Dam1-mNeonGreen molecule, we divided the median value of metaphase dot intensity (912) to 272 (16 kinetochores x 17 molecules per kinetochore). We assumed that the brilliance of one Dam1-mNeonGreen of 3.35 a.u. is similar to that of one molecule or Kar9-mNeonGreen. Thus, to calculate the amount of Kar9 molecules at the tip of the microtubule tip during metaphase we divided the mean value of the Kar9 dot intensity during metaphase (244 a.u.) by the calculated brilliance of one mNeonGreen molecule (3.35 a.u.), resulting in 72.9 Kar9 molecules at the tip of one astral microtubule.

For calculating the levels of over expressed Kar9, we multiplied the measured intensity of the focus with 2.5, the difference factor between the exposure times of over expressed Kar9 (100 ms) and of the endogenous or two copies of the gene (250 ms).

## KEY RESOURCES TABLE

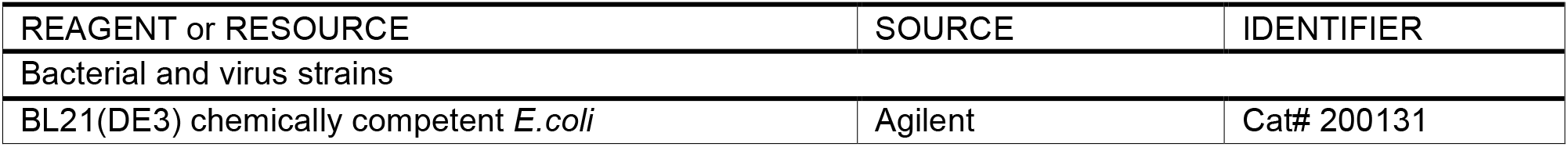
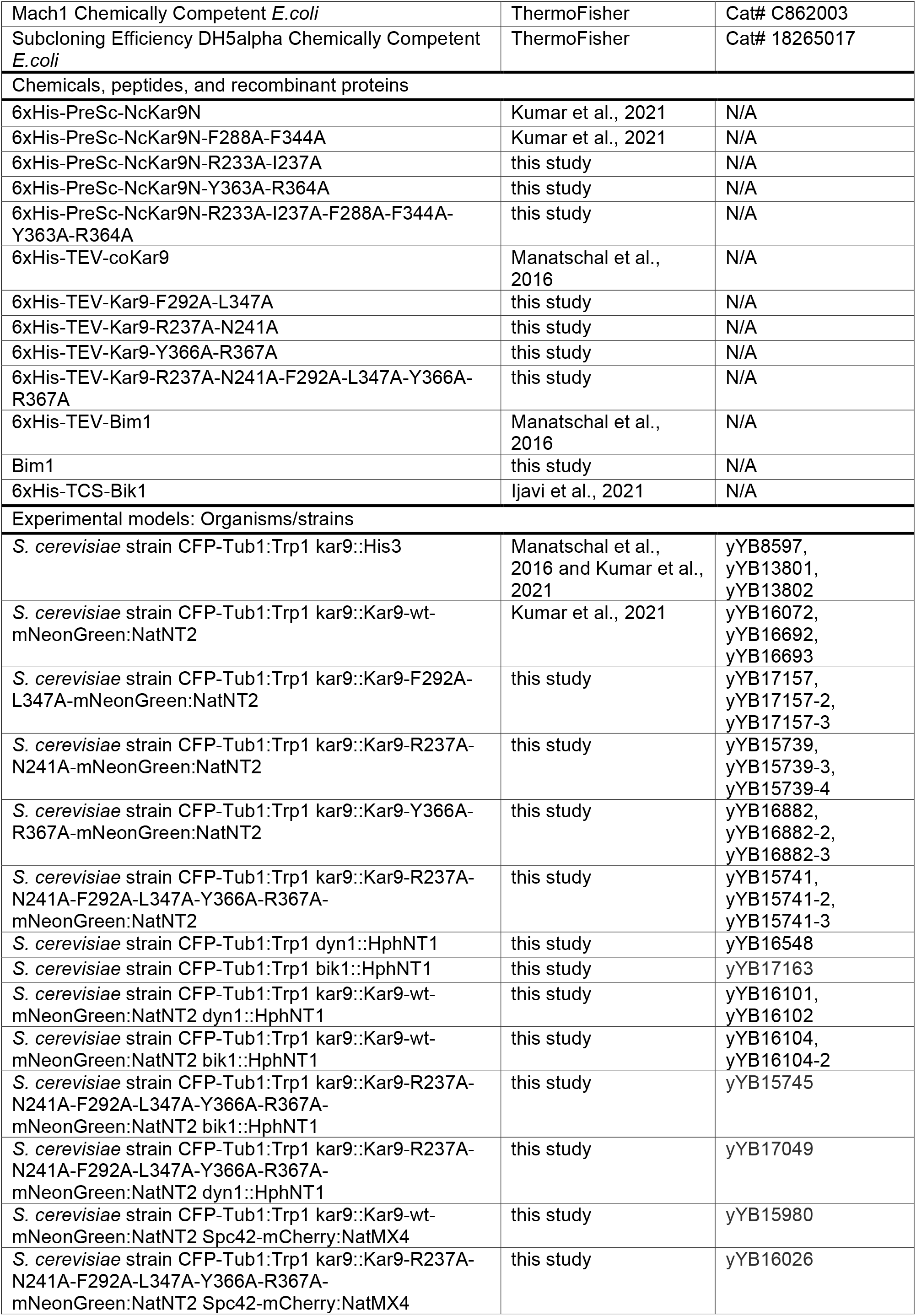
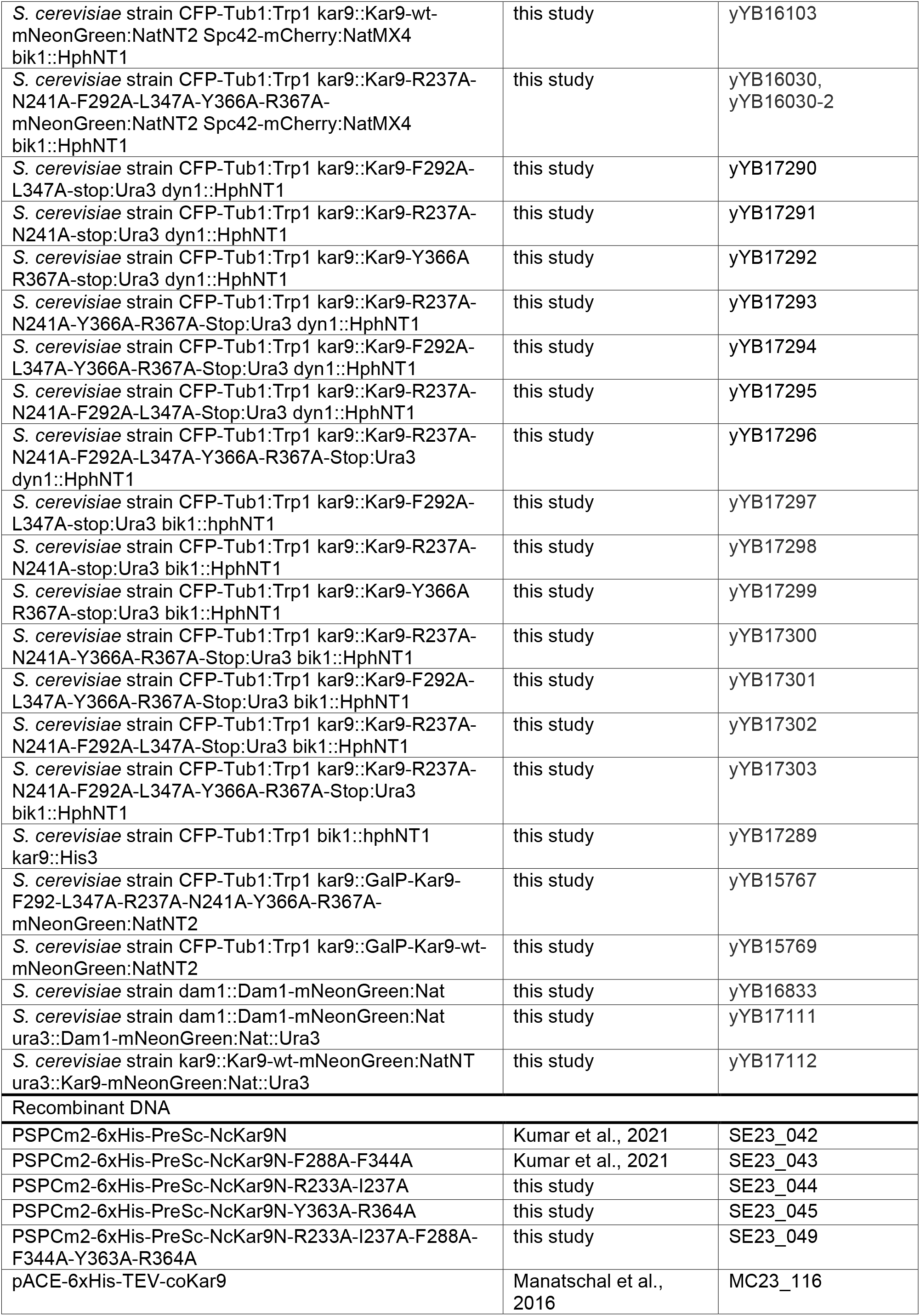
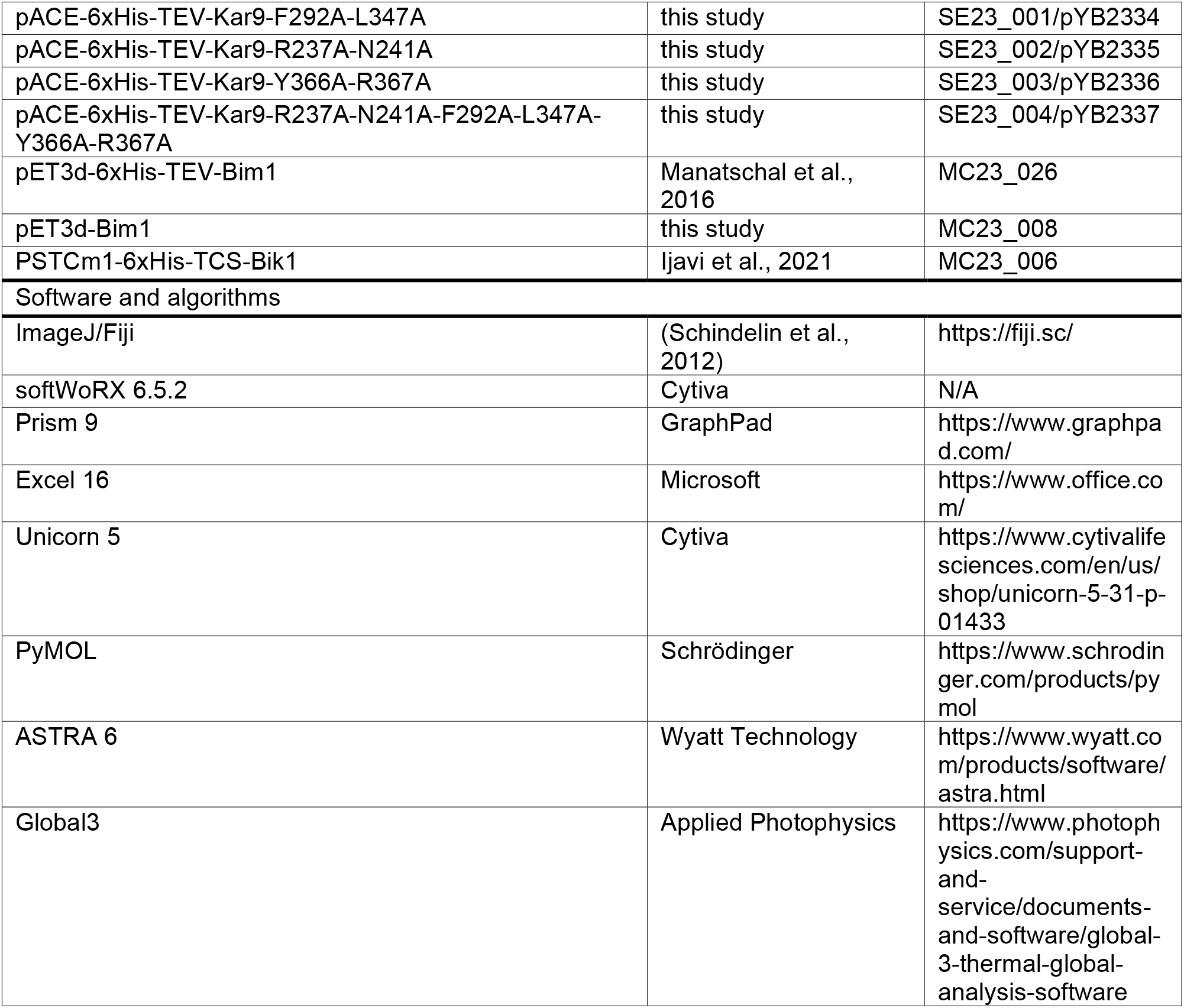

