## Supplementary Information for "High interaction valency ensures cohesion and persistence of a microtubule +TIP body at the plus-end of a single specialized microtubule in yeast"

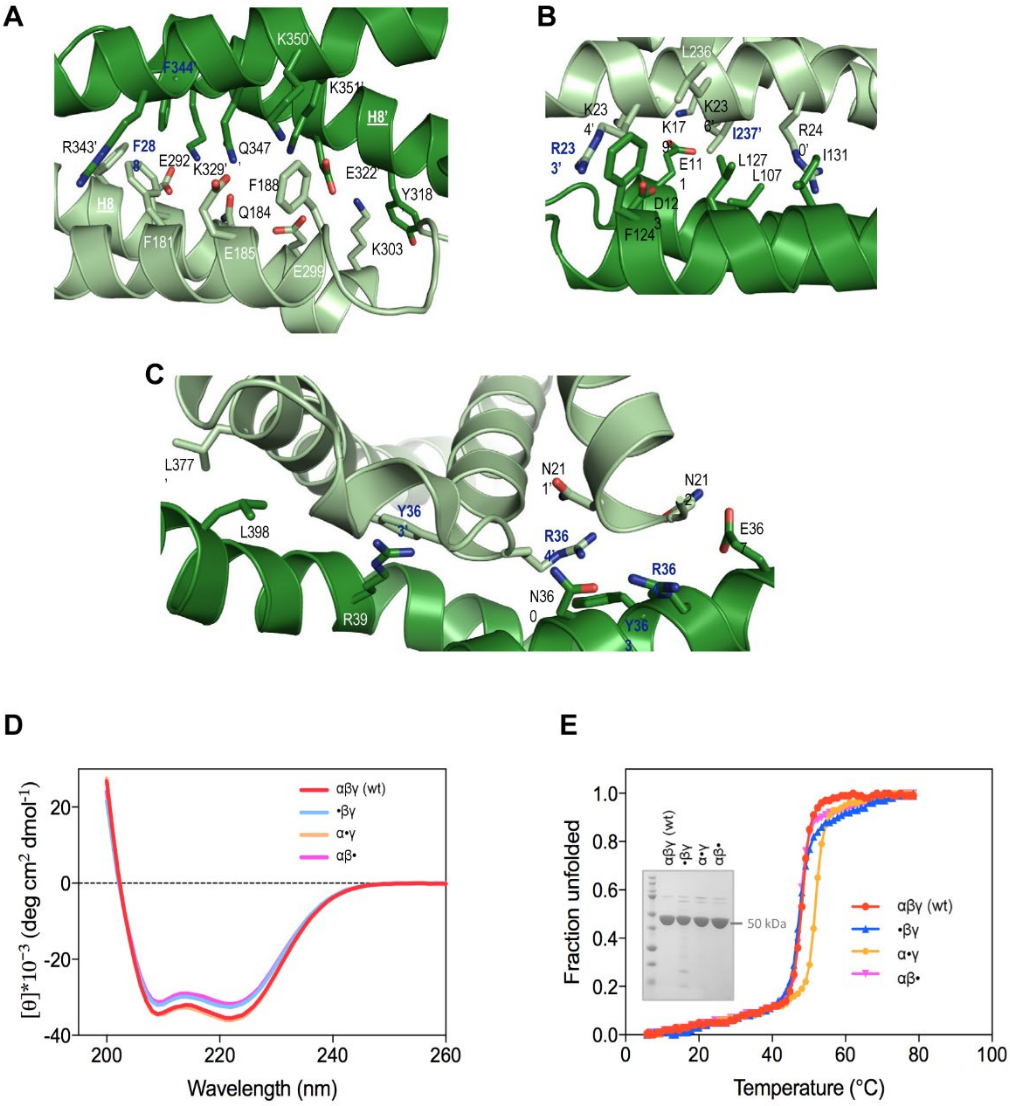


**Figure S1: Protomer contacts forming the NcKar9N crystal and NcKar9 dimer interface mutagenesis.**

**A-C**, Close up views of the $\alpha$ (A), $\beta$ (B), and $\gamma$ (C) interfaces seen between protomers in the NcKar9N crystal (PDB ID 7AG9). Interacting residue side chains are shown in stick representation. Oxygen and nitrogen atoms are colored in red and blue, respectively; carbon atoms are in dark green for chain A and in light green for chains B, C, and D. Residues that were mutated in the interfaces of the three crystallographic dimers are labeled in bold and blue lettering.

**D** and **E**, Far-ultra violet CD spectra (A) and thermal denaturation profiles (B) of His-NcKar9N-$\alpha\beta\gamma$ (red), His-NcKar9N-•$\beta\gamma$ (blue), His-NcKar9N-$\alpha$•$\gamma$ (yellow), and His-NcKar9N-$\alpha\beta$• (magenta). Inset in (B), SDS-PAGE analysis of the His-NcKar9N variants.

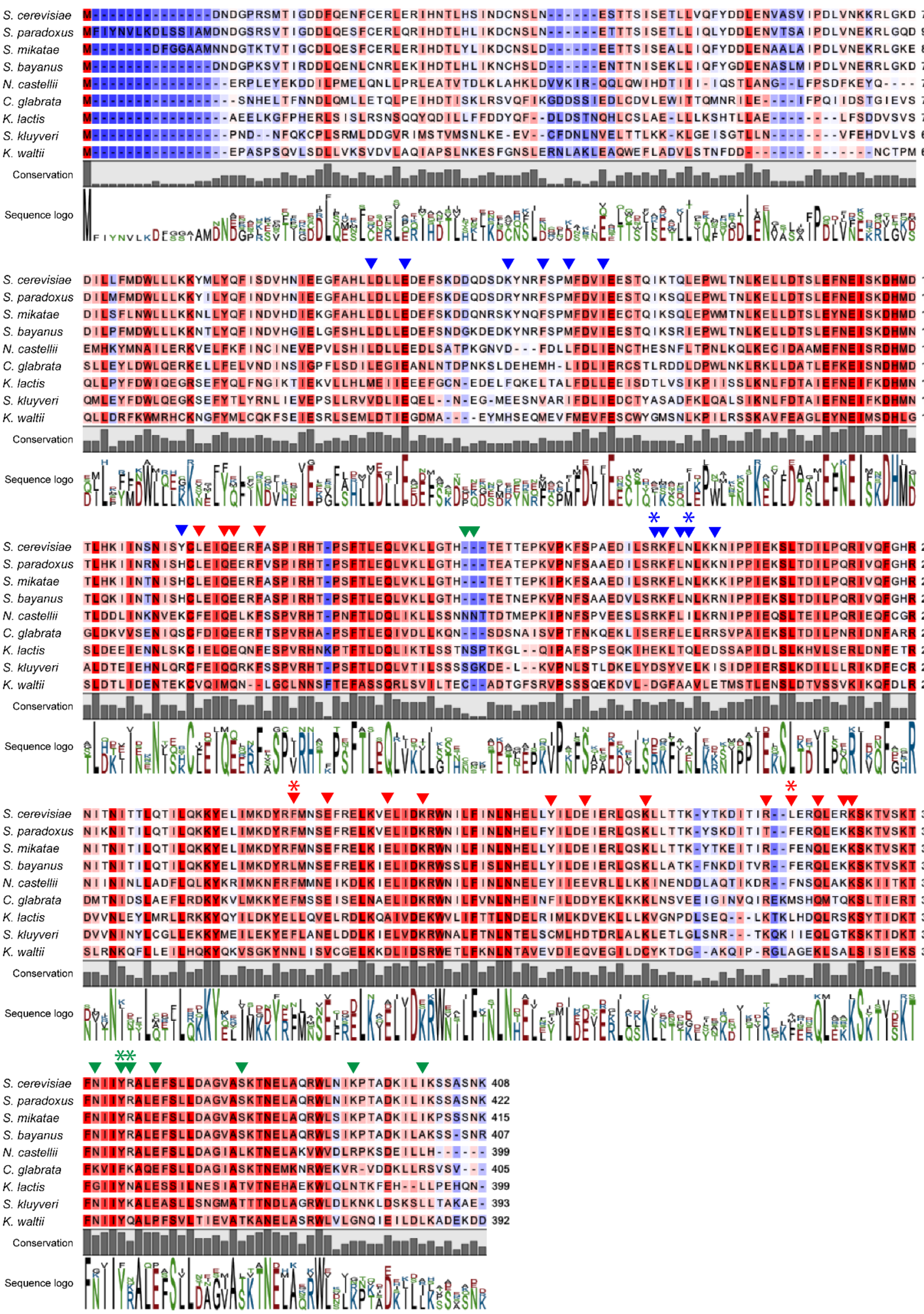


**Figure S2: Multiple sequence alignment of Kar9 homologues.**

Multiple sequence alignment of Kar9 homologues of closely and distantly related yeast species. Conserved amino acid residues are indicated by the background color from strong (red) to weak (blue) conservation, as well as by grey bars below the alignment. Consensus amino acids are indicated by the size of the letters below the alignment. Kar9 self-oligomerization interfaces are denoted by red (α), blue (β) and green (γ) arrowheads. Interface residues that were mutated in this study are indicated by asterisk in identical color.

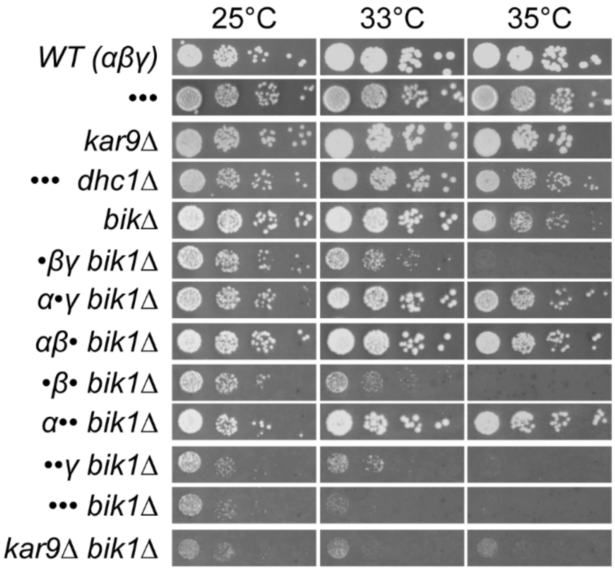


**Figure S3: Spot assay.**

Most relevant serial dilutions from the ones schematized in Figure 4A, tested at 25, 33, and 35°C.

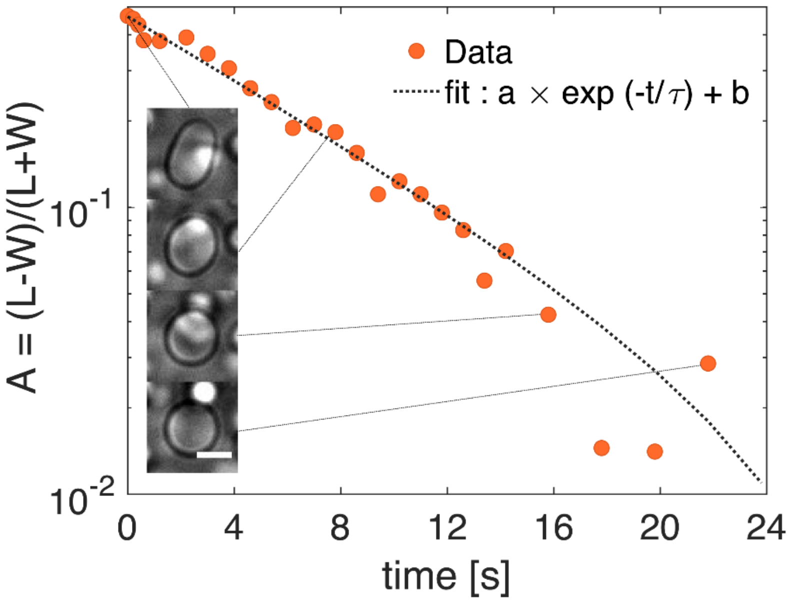


**Figure S4: Droplet fusion time quantification.**

Fusion of two Bik1 droplets at a later stage can be considered as a uni-axially deformed droplet reaching its final spherical state, driven by surface tension. Parameter A relates the major and minor axis of the droplet (L and W, respectively) and is plotted versus time, where an exponential fit allows to calculate the fusion time τ (see Materials and Methods, results are shown in Figure 1F). Scale bar in the inlet, 5 µm.
